# Genome size variation and species diversity in salamanders

**DOI:** 10.1101/065425

**Authors:** John Herrick, Bianca Sclavi

**Affiliations:** LBPA, UMR 8113, ENS Paris-Saclay, 95235 Cachan, France; Department of Physics, Simon Fraser University, 8888 University Drive, Burnaby, British Columbia VSA 1S6, Canada

**Keywords:** C-value, species diversity, body size, Urodela, geographical area

## Abstract

Salamanders (Urodela) have among the largest vertebrate genomes, ranging in size from 10 to 120 pg. Although changes in genome size often occur randomly and in the absence of selection pressure, non-random patterns of genome size variation are evident among specific vertebrate lineages. Several reports suggest a relationship between species richness and genome size, but the exact nature of that relationship remains unclear both within and across different taxonomic groups. Here we report i) a negative relationship between haploid genome size (C-value) and species richness at the family taxonomic level in salamander clades; ii) a correlation of C-value and species richness with clade crown-age but not with diversification rates; iii) strong associations between C-value and either geographical area or climatic niche rate. Finally, we report a relationship between C-value diversity and species diversity at both the family and genus level clades in urodeles.

## Introduction

Genome size in vertebrates varies more than three hundred fold from 0,4 picograms (pg) in pufferfish to over 120 picograms (pg) in lungfish (Gregory 2015). Most of the variation in vertebrate genome size corresponds to differences in non-coding DNA such as transposable elements, microsatellites and other types of repetitive and intergenic DNA (Metcalfe and Casane 2013). The DNA accounting for differences in genome size between related species has been considered devoid of any universal function such as gene regulation, structural maintenance and protection against mutagens (Palazzo and Gregory 2014). Genome size, however, is known to have a direct impact on important physiological parameters such as cell size, cell cycle duration and developmental time (Van’t Hof and Sparrow 1963, Cavalier-Smith 1978, Francis, Davies et al. 2008).

Numerous studies have reported either a positive or negative relationship between genome size (C-value) and species richness in vertebrates (Kraaijeveld 2010, Bromham 2011, Canapa, Barucca et al. 2015). C-value, for example, is positively correlated with species richness in fish (Mank and Avise 2006, Smith 2009), but negatively correlated in other taxa such as Amhibia and plants (Vinogradov 2003, Knight, Molinari et al. 2005). Likewise, across vertebrates, C-value has been found to be negatively associated with species richness, but only at the higher class-taxonomic level (Olmo 2006). The strongest association between genome size and species diversity was observed at C-values greater than 5 pg. Consistent with reduced species diversity in vertebrates with large genomes, extinction risk was found to increase with genome size (Vinogradov 2004). Consequently, species richness in a clade might correlate negatively with the proportion of repetitive DNA present in the respective species’ genomes (Olmo 2006), suggesting that either evolvability (propensity to speciate) or extinction risk varies according to the amount of non-coding DNA present in the genome.

Several hypotheses, both adaptive and non-adaptive, have been proposed to explain the relationship between C-value and species richness. The mutational hazard hypothesis, for example, proposes that large genomes impose a constraint on rates of speciation, presumably due to differences in life history traits such as body size and developmental rates (K-selected versus r-selected strategists) (Lynch and Conery 2003, Knight, Molinari et al. 2005, Larson 2011, Bromham, Hua et al. 2015). At the same time, larger genomes are more prone to DNA damage (Sparrow, Nauman et al. 1970, Conger and Clinton 1973, Vilenchik and Knudson 2003, Vilenchik and Knudson 2006). Consequently, larger genome sizes increase the number of mutations that can negatively impact genome stability and genetic integrity (Schneider, Liu et al. 2015, Mohlhenrich and Mueller 2016). The giant genomes of several plant and animal species might therefore have evolved either because of lower mutation rates in these taxa or simply because of genetic drift (Lynch and Conery 2003, Mohlhenrich and Mueller 2016, Lefébure, Morvan et al. 2017). These differing but not mutually exclusive hypotheses suggest that the relationship between C-value and species richness operates at several different levels of selection including the population level (Gregory 2004).

A more recent hypothesis based on the eukaryotic DNA replication and repair program proposes an underlying molecular mechanism to account for the frequently observed negative association between C-value and species richness (Herrick 2011). DNA repair systems are known to vary significantly among species, which might influence the respective rates of DNA sequence evolution (Britten 1986). Accordingly, as genomes increase in size during evolution, the DNA replication and repair program adapts to the growing mutational hazard, thereby limiting DNA damage to sub-lethal levels (Herrick and Bensimon 2008). Studies have shown, for example, that plant species with larger genomes repair damaged DNA at least as effectively as species with smaller genomes (Einset and Collins 2018), indicating that enhanced DNA repair systems compensate for the inevitably higher DNA damage levels incurred in larger genomes (Al Mamun, Albergante et al. 2016). Hence, instead of posing a hazard at the molecular levels of DNA damage and mutation rate, large genomes with disproportionate amounts of neutral or nearly neutral DNA might, paradoxically, act to stabilize the genome via enhanced DNA damage/repair systems (Herrick 2011). At the same time, correspondingly lower rates of genetic turnover (substitutions, recombination, insertions/deletions etc.) might limit the genetic diversity on which natural selection acts, thus limiting species diversity (Hidalgo, Pellicer et al. 2017).

Several studies support the DNA damage/repair hypothesis along three main lines of evidence. First, levels of genetic diversity in salamanders, frogs and mammals vary considerably in a genome size dependent manner (Pierce and Mitton 1980, Karlin and Means 1994). One study, for example, reported that Anura have higher levels of genetic polymorphism than Caudata but significantly smaller genomes (Nevo and Beiles 1991). Salamanders also have lower levels of heterozygosity in protein coding genes than do other vertebrates with smaller genome sizes (Matsui, Tominaga et al. 2008). Moreover, non-transforming salamanders, which have larger genomes on average, have correspondingly lower levels of heterozygosity compared to transforming salamanders (Shaffer and Breden 1989). Whether or not this variation is due to a genome size specific effect or some other explanation such as effective population size remains unresolved (Larson 1981, Parker and Kreitman 1982).

Second, two recent studies have shown that salamanders have on average exceptionally low nucleotide substitution rates (Herrick and Sclavi 2014, Mohlhenrich and Mueller 2016), confirming and extending earlier studies on mutation rates that suggest lower rates of molecular evolution in vertebrates with exceptionally large genomes (Dores, Sollars et al. 1999, Kozak, Costantino et al. 2005). The study on lungfish (C-value: 40 to 138 pg) showed, for example, that they have slower rates of molecular evolution than frogs (1 to 12 pg), while frogs have slower rates than mammals (1.7 to 6.3 pg). A more recent study on lungfish confirms this general trend, but with some exceptions (Biscotti, Gerdol et al. 2016).

Third, even earlier studies on rates of karyotype evolution showed that salamanders have slower rates of genome evolution than frogs, while frogs have slower rates than mammals (Wilson, Sarich et al. 1974, Bush, Case et al. 1977, Bengtsson 1980). A more recent study on karyotypic diversification rates at the family level of mammalian clades supports these findings (Martinez, Jacobina et al. 2017). The observed slower karyotype diversification rates in large genomes are consistent with other observations that large genomes in plants, animals and insects experience slower rates of DNA loss and, consequently, genome size evolution (Wilson, Sarich et al. 1974, Bensasson, Petrov et al. 2001, Sun, López Arriaza et al. 2012, Kelly, Renny-Byfield et al. 2015, Pellicer, Hidalgo et al. 2018).

Together, these findings suggest that large genomes tend to be associated with slower rates of molecular evolution (genetic and genomic turnover), which might be reflected in terms of species diversity within and across different but closely related taxonomic groups and clades (Böhne, Brunet et al. 2008). The Anura, for example, have smaller genomes on average that are evolving more rapidly compared to Caudata. (Liedtke, Gower et al. 2018). At the same time, the Anura contain correspondingly more species-rich clades than do Caudata (Pyron and Wiens 2011). This observation raises the question of whether an inverse relationship between species richness and genome size, and/or genome diversification, persists at lower taxonomic levels within the salamander clade as the earlier studies on rates of karyotype evolution have suggested (Bush, Case et al. 1977, Bengtsson 1980).

Genomic turnover (eg. substitutions, translocations, deletions and amplifications, etc.) is more easily assessed in terms of variation in genome size across taxonomic clades with large genomes, since observable differences in genome size reflect underlying mutational processes occurring over evolutionary time (Feschotte and Pritham 2007, Kapusta, Suh et al. 2017). The large and widely differing sizes of salamander genomes therefore offer an attractive proxy to examine any potential association between species richness and genome size variation (Sessions 2008). Several variables likely influence the dynamics of both species richness and genome size, in particular evolutionary time (as measured by clade crown age), geographic range and environmental heterogeneity. How these variables impact genome size evolution has been the focus of intense interest for many years. In the following, we examine the relationships between species richness, C-value and genome size diversity within the Caudata.

## Materials and methods

### Genome sizes

were obtained from the Animal Genome Size Database (Gregory 2015). C-value refers to haploid nuclear DNA content (1C). Reported polyploids, when indicated in the Animal Genome Size Database, were removed from the analyses. Average C-values were determined for each species when more than one C-value is recorded in the database. These values were used to calculate the average C-value of each family-level clade and each genus-level clade. The distributions in genome size for each salamander family and among the genera of Plethodontidae have been published previously (Herrick and Sclavi 2014).

### The data on crown age, stem age, species diversity, niche rate and geographic area

were obtained from Pyron and Wiens (Pyron and Wiens 2013). Both the maximum likelihood (ML) and the time-calibrated trees used here were obtained from those of Pyron and Wiens (Pyron and Wiens 2011, Pyron 2011). They obtained the time-calibrated tree by determining divergence times from a set of fossil constraints using treePL developed by S.A. Smith (Smith and O’Meara 2012), and applied to the ML phylogeny determined previously for 2871 species using data from 3 mitochondrial and 9 nuclear genes (Pyron and Wiens 2011). They determined species diversity from the assignment of all known amphibian species to clades as classified in their phylogeny (Pyron and Wiens 2011). Species richness at the genus level was determined by counting the number of species per genus in the Pyrons and Wiens tree. Niche rate is defined as in Kozak and Wiens (Kozak and Wiens 2010). The radiation time at the genus level was obtained from TimeTree (http://www.timetree.org). The species level area was obtained from the IUCN Red List (http://www.iucnredlist.org/initiatives/amphibians/analysis/geographic-patterns). The SAGA software (Conrad *et al* 2015) was used to measure the genus-level area.

The Pyrons and Wiens ML and time-calibrated trees were used to create trees at the family or genus level. Species were assigned to families at first using the *taxize* package in R (Chamberlain and Szöcs 2013). These were manually verified against the taxonomy of the Pyrons and Wiens tree. The HighLevelTree function in the *EvobiR* package in R by Heath Blackmon was used to obtain the family or genus level trees (evobiR: evolutionary biology in R. R package version 1.0. http://CRAN.R-project.org/package=evobiR).

### Regression analysis

The univariate and multivariate pgls analysis was carried out in R with the *caper* package. The maximum likelihood value of lambda was allowed to vary while kappa and delta were set to 1 as in Kozak and Wiens (Ecology and Evolution 2016). For the pgls analysis we used the time-calibrated family or genus tree obtained from the Amphibia tree of Pyron and Wiens (Pyron and Wiens 2013) as described above, or the phylogenetic tree from Kozak and Weins (Kozak and Wiens 2010) for the Plethodontidae dataset. Data used in the regression analysis involving the coefficient of variation of C-value were arcsine square root transformed and assessed using the Shapiro-Wilks test for normality (see supplementary material). The Benjamini-Hochberg procedure was used to rank p-values with a False Discovery Rate (FDR) of 0.05.

## Results

### Species diversity and C-value variation at the family-level of salamander clades

The oldest known urodeles date from 166 to 168 Mya (Marjanovic and Laurin 2014, Laurin, Canoville et al. 2015). The urodeles inhabit a wide variety of ecological niches and exhibit a large diversity of life-history traits, including small and large body sizes, paedomorphy, metamorphosis and direct development (Wake 2009). In an earlier study of Amphibia, Pyron and Wiens revealed a number of ecological correlates between species diversity and variables such as geographical latitude, geographic range and environmental energy (Pyron and Wiens 2013). Species diversity in frogs, salamanders and caecilians also varies according to abiotic factors such as humidity and temperature and biotic factors such as productivity and rates of diversification (extinction and speciation). Figure 1 shows the family level phylogenetic tree while Figure 2 shows the genus level phylogenetic tree, both derived from Pyron and Wiens (Pyron and Wiens 2011), that were used here to investigate the relationship between salamander genome size and species diversity. Substantial variation in species diversity, body size and C-value among and within clades at the family and genus level is apparent in the phylogenetic tree of the Urodela.

**Figure 1.**
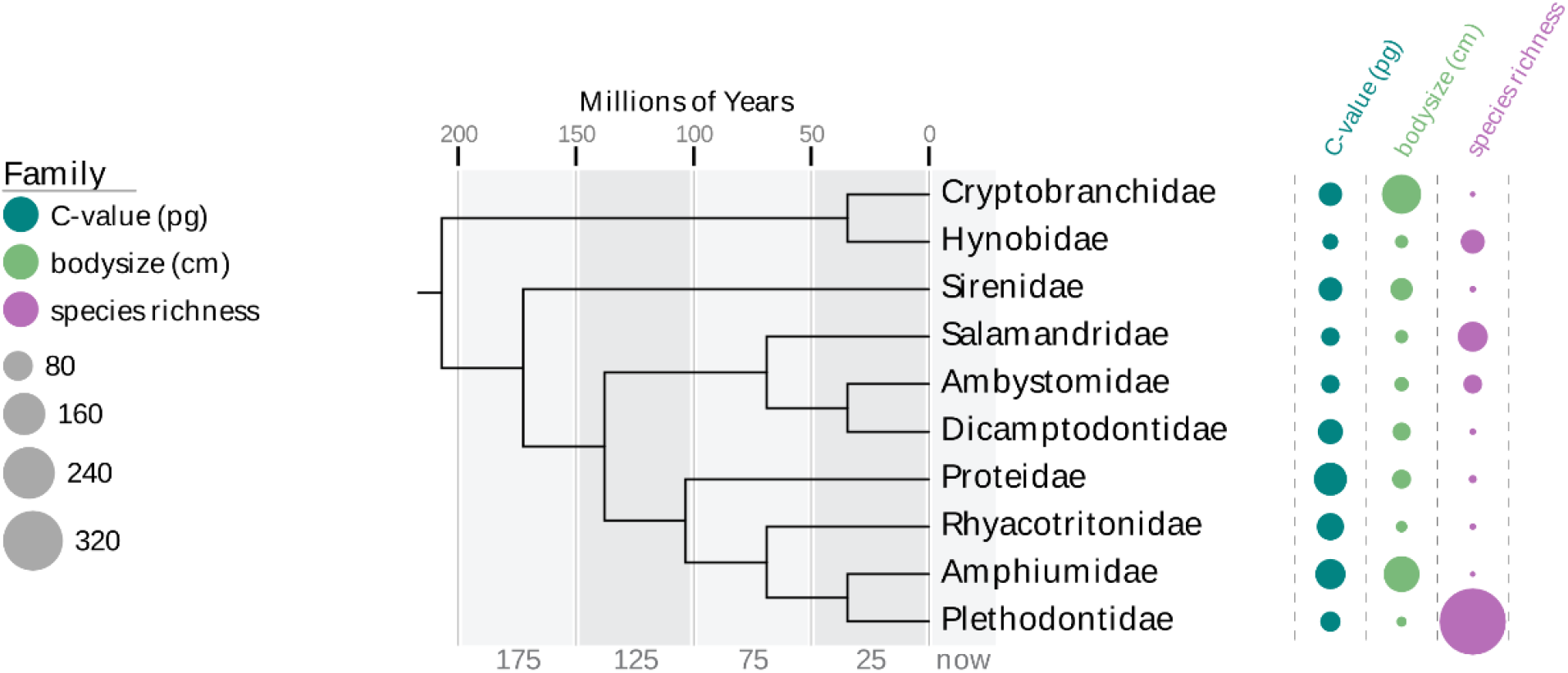
Time-calibrated phylogenetic tree of Caudata families derived from Pyron and Wiens 2011 (Pyron and Wiens 2011). Figure created using EvolView (Zhang, Gao et al. 2012).

**Figure 2.**
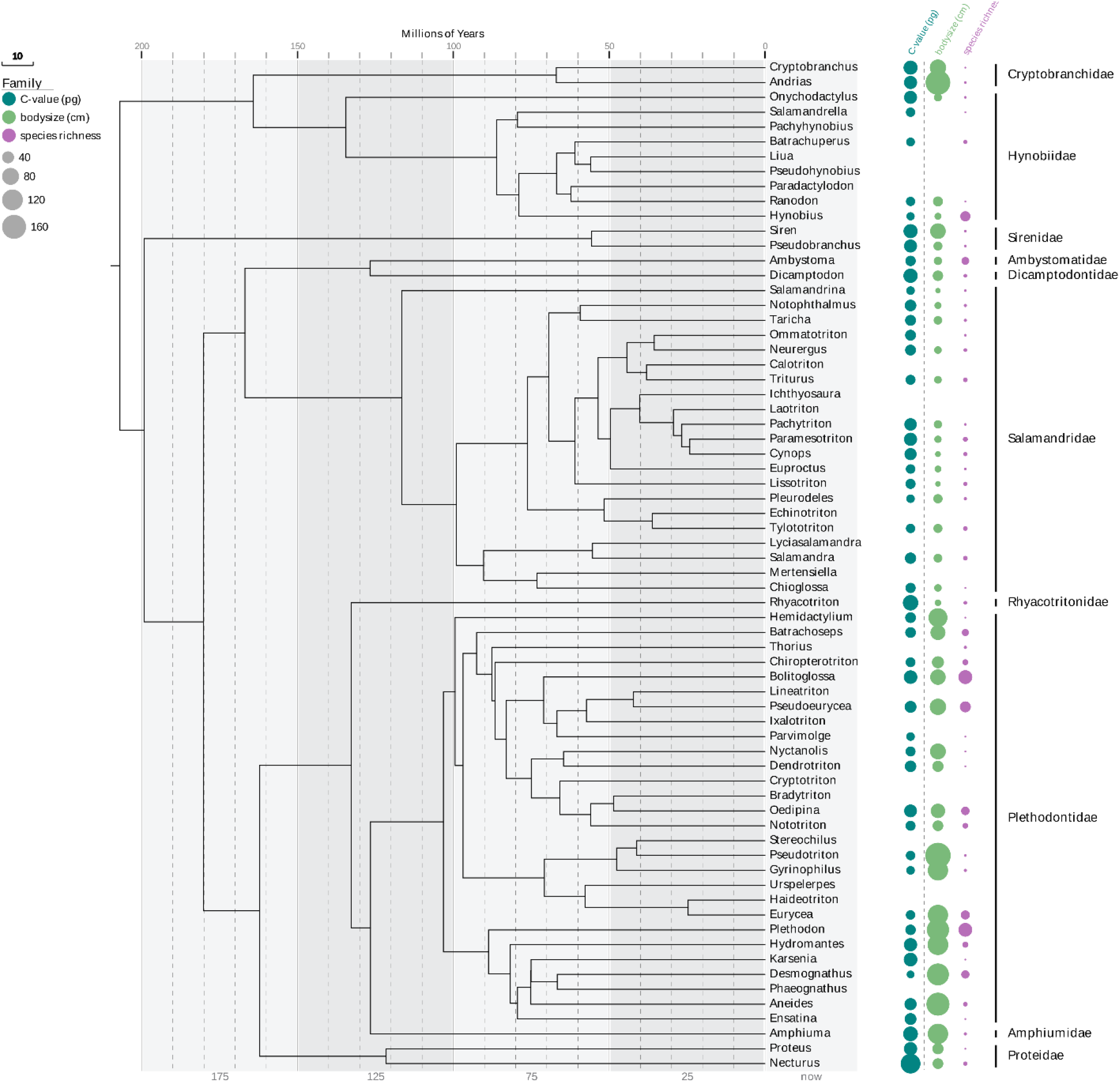
Time-calibrated phylogenetic tree of Caudata genera derived from Pyron and Wiens 2011 (Pyron and Wiens 2011). Figure created using EvolView (Zhang, Gao et al. 2012).

### Crown age, but not diversification rate, correlates with clade diversity

Among animal taxa it has been reported that clade age rather than diversification rate explains species richness (McPeek and Brown 2007). A more recent study of amphibians, birds and mammals supports the finding that time (older lineages), and not diversification rates, explains extant species richness (Marin and Blair Hedges 2016). A significant positive correlation between crown group age at the family level and species diversity has also been reported in salamanders (Eastman and Storfer 2011). In contrast to these reports, our initial phylogenetic generalized least squares (PGLS) analysis at the family-level of Urodela did not confirm a significant relationship between crown age and species richness.

We determined, however, in preliminary OLS regression analyses that the family Proteidae represents an outlier (studentized residual: 2.35; studentized deleted residual: 3.94). We further determined that the outlier status of the Proteidae is due to a single monotypic genus: the European *Proteus anguinus*, the only genus in the Proteidae clade found outside of North America. Monotypic groups complicate estimates of crown age (Wiens 2017), which might influence the regression analyses performed here. Excluding this monotypic genus from the regression analysis changes the Proteidae family-level crown age of 121 Mya to that of the Necturus genus-level crown age of 13 Mya (www.timetree.org).

We therefore conducted the following PGLS analyses both *with and without* the single species *Proteus anguinus* to test the effect this species has on the association between the investigated variables (see Table S1 and Table 1). Phylogenetic generalized least squares (PGLS) analysis confirmed that species diversity in salamanders increases with clade crown age: older clades tend to have higher species diversity than comparatively younger clades, as expected (Table 1). In contrast to crown age, we found no correlation between stem age and clade diversity at the family level (R^2^ = 0.1; P = 0.8), in agreement with earlier reports on other taxa (Rabosky, Slater et al. 2012, Stadler, Rabosky et al. 2014, Wiens 2017).

**Table 1.**
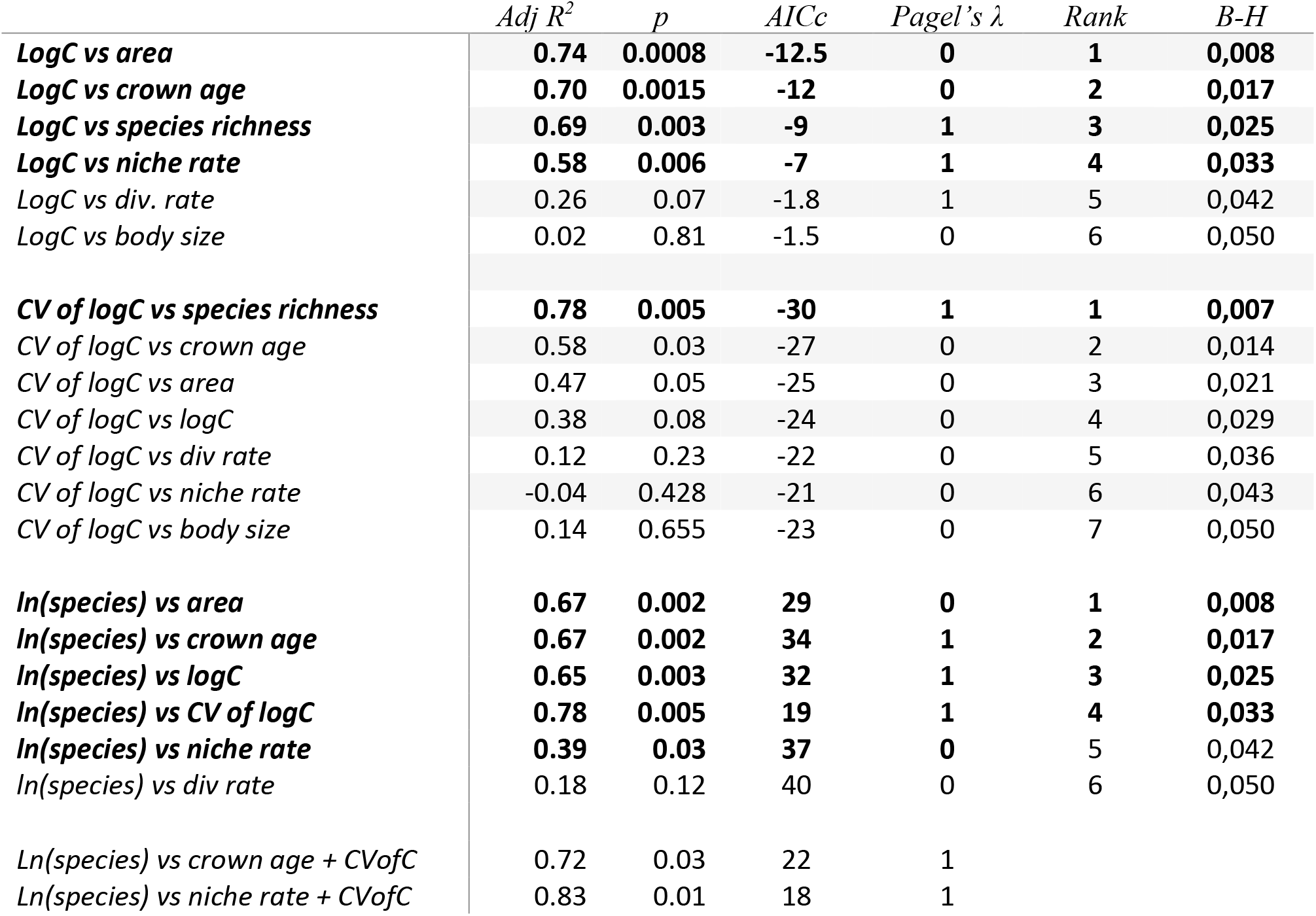
PGLS analysis for Salamander family-level clades of correlations of C-values with data from Pyrons and Wiens (Pyron and Wiens 2013) without *Proteus anguinus*. The Benjamini-Hochberg procedure was used with a False Discovery Rate of 0.05. The results of multiple regression improving the AIC score are shown.

### Average C-value correlates with crown age but not diversification rate

The ancestral Amphibia genome (C-value ~ 3 pg) has experienced massive amplification during the evolution of the Urodela (Organ, Canoville et al. 2011, Organ, Struble et al. 2016). An early ancestor of salamanders, for example, has been estimated to have a large genome size of approximately 33.1 pg, which is similar to the reconstructed ancestral genome size of ~32 pg (Sessions 2008, Laurin, Canoville et al. 2015). An earlier study suggested that genome size in salamanders has increased with time, although the exact rate of increase remains unclear (Martin and Gordon 1995, Sessions 2008). We therefore examined the relationship between phylogenetic stem age and genome size but found no significant relationship between stem age and C-value (not shown). Examination of the relationship between crown age and C-value using the Pyron and Wiens dataset, in contrast, revealed a significant negative relationship between C-value and crown age (R^2^ = 0.7 P = 0.0015; Table 1).

### C-value, but not body size, is associated with species richness

Figure 1 suggests that species diversity is negatively associated with both C-value and body size. Our analysis revealed an inverse relationship between species richness and body size; however, the correlation was not significant (Table 1). The relationship between species richness and C-value, in contrast, is strongly negative and significant at the family taxonomic level (R^2^ = 0.69 P = 0.003; Table 1). This negative trend is readily apparent in the phylogenetic tree: the sister clades Hynobidae and Cryptobranchidae, for example, differ over 2X with respect to average genome size and over 10X with respect to species diversity (Figure 1). The other two sister clades in Figure 1 exhibit a similar trend (Amphiumidae: Plethodontidae; Dicamptodontidae: Ambystomatidae).

### C-value, species richness, geographic area and climatic niche rate

Older clades (crown age) have had more time to disperse over larger areas, suggesting that geographic area might correlate positively with species richness. PGLS analysis revealed a strong relationship between family clade diversity and geographic area (R^2^ = 0.67, P = 0.002; Table S1). Likewise, a strong negative correlation was found between C-value and geographic area: clades with smaller average genome sizes occupy larger geographic areas (R^2^ = 0.74; P = 0.0008; Table 1).

In order to better understand the relationship between C-value and geographic area we next examined the relationship between C-value and niche rate. Since larger geographic areas are expected to comprise higher levels of both habitat and niche diversity, we examined the individual relationships between niche rate, species richness and C-value. PGLS analyses revealed a negative relationship between niche rate and C-value (Table 1). Positive relationships were found between area and species richness and between niche rate and area (Table S2). Additionally, our analysis revealed that niche rate and diversification rate are related at the family-level of Urodela clades (R^2^ = 0.44; P = 0.02; Table S2), as previously reported for lower taxonomic levels (Kozak and Wiens 2010). This observation is consistent with a more extensive study showing a strong correlation between diversification rate and niche rate at family level clades in mammals (Castro-Insua, Gómez-Rodríguez et al. 2018).

### Genome size diversity and species diversity

Together, our results suggest a potential relationship between species diversity and genome size diversity. We next investigated if clades with higher levels of species diversity have correspondingly higher levels of genome size diversity. Genome size is expected to evolve in a manner that is proportional to C-value, suggesting that larger genomes are changing faster in size compared to smaller genomes (Oliver, Petrov et al. 2007). The coefficient of variation (CV) of the log of C-value was used here to assess genome size diversity (see materials and methods). Comparing salamander families, there appeared to be a negative relationship between average C-value in each clade and its corresponding CV; however, the relationship was not significant (R^2^ = 0.38; P = 0.08; Table 1).

Examining the relationships between CV of C-value and the other investigated variables, a strong relationship was found between genome size diversity and species richness at the family taxonomic level (Table 1), as expected if genome size has been evolving in parallel with species richness since the emergence of the different family-level salamander clades. Significant correlations between CV of C-value and either crown age or area were also found; however, only the correlations between species richness and CV of C-value remained significant following Benjamini-Hochberg analysis (Table 1).

### Clade age, species richness and C-value at the genus level

Extending our examination of the relationship between clade age and species richness to genus-level clades across the Urodela phylogenetic tree, we initially failed to find a clear correlation between clade age and species richness in 50 different genera. Excluding the monotypic phyla, however, revealed a significant correlation between genus radiation time and species richness (R^2^ = 0.62; P = 2 x 1E-9, Table 2). When examining the relationship between CV of C-value and radiation time we also found a significant correlation between these two variables (R^2^= 0.21; P = 0.008; Table 2). Multiple regression analysis showed that the contribution of each variable, CV of C or radiation time, to species richness was not substantially different from their individual univariate contributions (Table 2). Phylogenetic path analysis at the genus level further suggests that the best of all possible models corresponds to the one in which species diversity and genome size diversity depend similarly on radiation time, as expected (not shown).

**Table 2.**
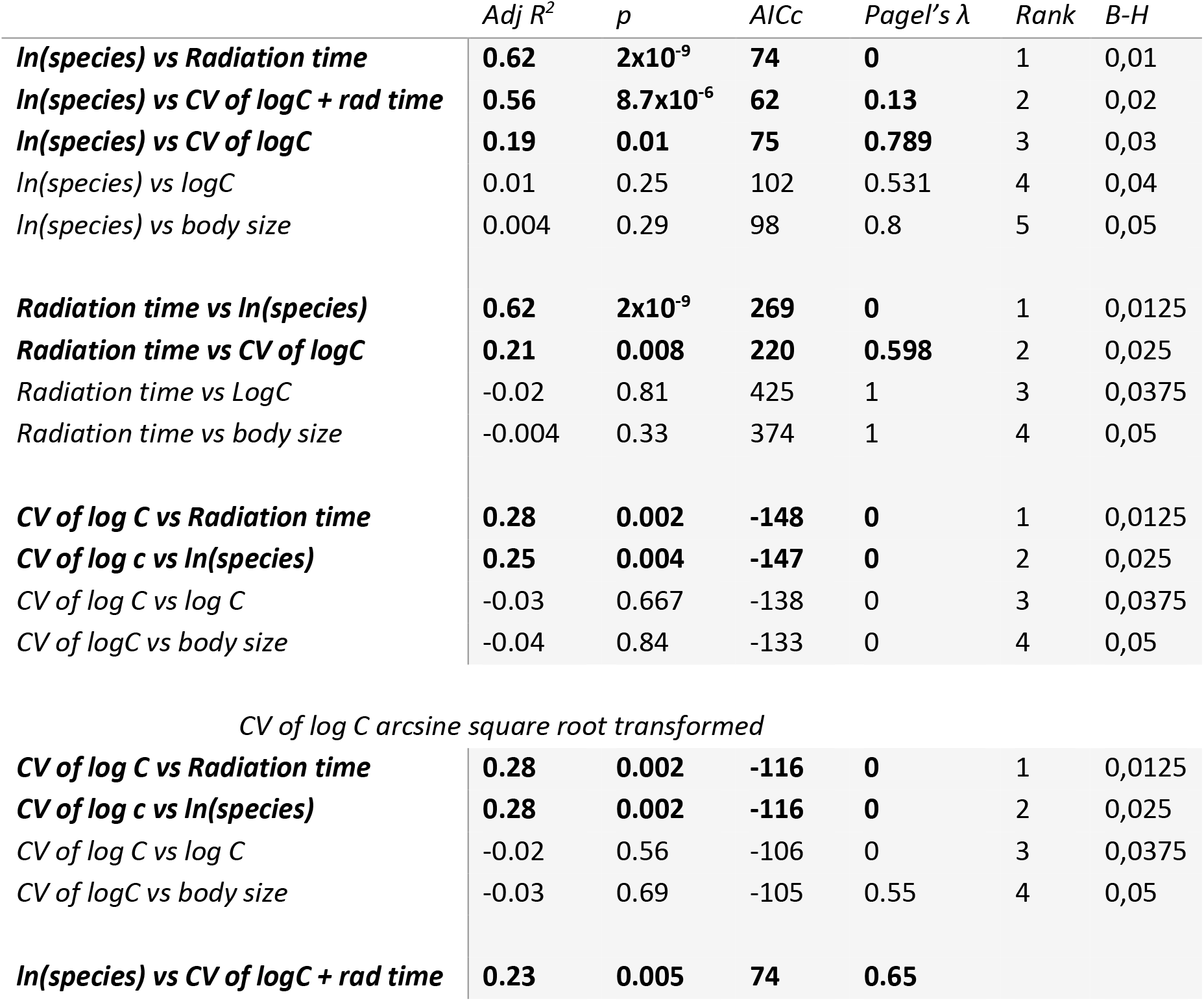
Results of the pgls analysis at the genus level. The analysis with species richness was carried out in the absence of the monophyletic genera.

A significant relationship has previously been reported between diversification rate and niche rate in 15 plethodontid clades (Kozak and Wiens 2010). Using the values of niche rate for plethodontids reported in Kozak and Wiens 2016, we found a significant association between C-value and niche rate at this lower taxonomic level (R^2^ = 0.48; P = 0.002; Table 3), similar to what was observed at the family level (R^2^ = 0.58; P = 0.006; Table 1). In contrast, we did not find that C-value or CV of C-value is associated with either crown age, geographic area, diversification rate, or species diversity in the plethodontid clade (Table 3).

**Table 3.** PGLS analysis of Plethodontidae data from Kozak and Wiens (Kozak and Wiens 2016). The Benjamini-Hochberg procedure was used with a False Discovery Rate of 0.05.

|  | *Adj R<sup>2</sup>* | *p* | *AICc* | *Pagel's λ* | *Rank* | *B-H* |
| --- | --- | --- | --- | --- | --- | --- |
| ***LogC vs niche rate*** | **0.48** | **0.002** | **-19** | **0** | **1** | **0,01** |
| *LogC vs area* | 0.14 | 0.09 | -12.65 | 0.313 | 2 | 0,02 |
| *LogC vs div. rate* | -0.02 | 0.45 | -11 | 1 | 3 | 0,03 |
| *LogC vs crown age* | -0.07 | 0.76 | -11 | 1 | 4 | 0,04 |
| *LogC vs species richness* | -0.07 | 0.8 | -11 | 1 | 5 | 0,05 |

## Discussion

We report here a relationship between average genome size and species richness at the family level of Urodela clades (Table 1). At both genus and family levels, a significant relationship was also found between C-value diversity (CV of C) and species richness in a clade. Phylogenetic path analysis at the genus level suggests that these two variables, genome size diversity and species richness, depend similarly on crown age, suggesting that these two traits are evolving independently of each other (not shown). Together, our findings provide evidence that C-value and variation in C-value constitute additional traits associated with species diversity, at least in urodeles. Examining other factors such as clade age, geographic area and rate of climatic-niche evolution suggests that the relationship between genome size and species diversity is mediated directly or indirectly through several different variables.

What is the nature of the relationship between average C-value and these different ecological variables in the Urodela? Sessions reported that the range in genome size in family-level Urodela clades tends to increase as their average C-values decrease (Sessions 2008), suggesting a potential relationship between genome size diversity and average genome size in a clade. Our analysis does not support a significant relationship between CV of genome size and C-value. We did find, however, a significant relationship between CV of genome size and species richness, suggesting that diversification of genome size coincided with diversification of species in a clade (Table 1). In contrast, genome size diversity (CV of C) is not significantly related to either niche rate, geographic area or diversification rate at the family taxonomic level (Table 1). These observations suggest that changes in genome size are associated directly or indirectly with speciation events in urodeles, consistent with findings in plants that rates of genome size evolution correlate with rates of speciation (Puttick, Clark et al. 2015).

We note that the observations made here concerning average C-value and species diversity apply predominantly to the family-level of Urodela clades. At lower taxonomic levels such as the plethodontids, the relationship between genome size and species richness shows no consistent pattern. The Bolittoglossinae, for example, tend to have larger clade-average genome sizes but higher species diversity than other genera in the plethodontids. Indeed, the Plethodontidae as a group exhibit the highest levels of species diversity among family-level Urodela clades (Wake 2009), and have correspondingly elevated levels of genome size diversity and substitution rates (Herrick and Sclavi 2014). We suggest that genome size diversity, rather than genome size itself, reflects the correspondingly higher substitution rates previously reported for the Plethodontidae (Herrick and Sclavi 2014).

Together, our analyses support the *Geographic Range Hypothesis*, according to which taxa with wider geographic distributions have higher probabilities that genetic changes such as indels, chromosomal inversions and transposon-mediated modifications in genome size will become fixed in a population (Feder, Gejji et al. 2011, Martinez, Jacobina et al. 2017). If habitat and niche availability both increase with geographic area, then our observations suggest that changes in genome size in urodeles might have occurred *in parallel* with adaptations that made available habitats and niches more accessible to dispersing ancestral populations, but only over longer evolutionary periods (family level clades).

How might variation in C-value contribute to speciation events? The recent study on karyotypic diversification rates at the family level of mammalian clades suggests that large, past geographic distributions in heterogenous environments might have favored higher levels of chromosomal diversity; or conversely, higher rates of chromosomal diversification might have promoted colonization of new habitats and expanding geographic ranges (Martinez, Jacobina et al. 2017). These results are consistent with the earlier observations on the rate of karyotype diversification in mammals, frogs and salamanders (Wilson, Sarich et al. 1974, Wilson, Bush et al. 1975, Bush, Case et al. 1977, Bengtsson 1980), suggesting that taxonomic groups with larger genomes on average have slower rates of genome evolution (Hooper and Price 2015, Leaché, Banbury et al. 2016).

Based on these findings, we propose that rates of variation in C-value and genomic organization (heterochromatin content and gene synteny), in addition to changes in genome size, might coincide with rates of variation in species richness. These observations suggest that changes in C-value, and hence changes in the amount and/or organization of non-coding DNA in the vertebrate genome, are associated with the allelic incompatibility believed to drive reproductive isolation and speciation in salamanders. We are currently investigating the hypothesis that genome diversification rates and corresponding levels of genome size and karyotype diversity, rather than absolute C-value, explain rates of molecular evolution and rates of speciation in Amphibia and other eukaryotes.

## Acknowledgements

BS is supported by a grant from Human Frontier Science Program (RGY0079). JH benefited from support from John Bechhoefer’s lab, Physics Department, Simon Fraser University.

## Supplementary Information

**Supplementary Table 1.** PGLS analysis for Salamander family-level clades of correlations of C-values with data from Pyrons and Wiens (Pyron and Wiens 2013) including *Proteus anguinus*.

|  | *Adj R<sup>2</sup>* | *p* | *AICc* | *Pagel's λ* |
| --- | --- | --- | --- | --- |
| ***Area vs logC*** | **0.77** | **0.0005** | **329** | **0** |
| ***Area vs species richness*** | **0.69** | **0.002** | **332** | **0** |
| ***Area vs niche rate*** | *No change* |  |  |  |
| *Area vs CV of logC* | 0.27 | 0.13 | 238 | 0 |
| ***Area vs crown age*** | **0.45** | **0.02** | **337** | **0** |
| ***LogC vs area*** | **0.80** | **0.0003** | **-17** | **1** |
| ***LogC vs species richness*** | **0.69** | **0.002** | **-13** | **1** |
| *LogC vs sr/crown* | 0.12 | 0.17 | -4 | 0 |
| ***LogC vs niche rate*** | **0.60** | **0.005** | **-11** | **1** |
| *LogC vs crown age* | 0.31 | 0.056 |  | 0.001 |
| *LogC vs div. rate* | 0.22 | 0.09 | -4 | 1 |
| *CV of logC vs area* | 0.27 | 0.13 | -26 | 0 |
| *CV of logC vs logC* | 0.07 | 0.28 | -25 | 0 |
| *CV of logC vs niche rate* | -0.14 | 0.62 |  | 0 |
| ***CV of logC vs crown age*** | **0.7** | **0.01** | **-32** | **0** |
| ***CV of logC vs species richness*** | **0.49** | **0.05** | **-29** | **0** |
| *CV of logC vs div rate* | -0.05 | 0.44 | -24 | 0 |
| ***Species richness vs niche rate*** | **0.38** | **0.03** | **35** | **0** |
| *Species richness vs div rate* | 0.18 | 0.12 | 38 | 0 |
| ***Species richness vs crown age*** | **0.36** | **0.04** | **36** | **0.001** |

**Supplementary Table 2.** PGLS analysis for Salamander family-level clades with data from Pyrons and Wiens (Pyron and Wiens 2013) excluding *Proteus anguinus*.

|  | *Adj R<sup>2</sup>* | *p* | *AICc* | *Pagel's λ* |
| --- | --- | --- | --- | --- |
| ***Niche rate vs area*** | **0.60** | **0.005** | **10.5** | **0.8** |
| ***Niche rate vs div rate*** | **0.44** | **0.02** | **13** | **0** |
| ***Niche rate vs species richness*** | **0.39** | **0.03** | **14** | **0** |

## References

Bengtsson, B. O. (1980). “Rates of karyotype evolution in placental mammals.” Hereditas 92(1): 37-47.

Bensasson, D., D. A. Petrov, D. X. Zhang, D. L. Hartl and G. M. Hewitt (2001). “Genomic gigantism: DNA loss is slow in mountain grasshoppers.” Mol Biol Evol 18(2): 246-253.

Biscotti, M. A., M. Gerdol, A. Canapa, M. Forconi, E. Olmo, A. Pallavicini, M. Barucca and M. Schartl (2016). “The Lungfish Transcriptome: A Glimpse into Molecular Evolution Events at the Transition from Water to Land.” Sci Rep 6: 21571.

Britten, R. J. (1986). “Rates of DNA sequence evolution differ between taxonomic groups.” Science 231(4744): 1393-1398.

Bromham, L. (2011). “The genome as a life-history character: why rate of molecular evolution varies between mammal species.” Philosophical transactions of the Royal Society of London. Series B, Biological sciences 366(1577): 2503--2513.

Bromham, L., X. Hua, R. Lanfear and P. F. Cowman (2015). “Exploring the Relationships between Mutation Rates, Life History, Genome Size, Environment, and Species Richness in Flowering Plants.” Am Nat 185(4): 507-524.

Bush, G. L., S. M. Case, A. C. Wilson and J. L. Patton (1977). “Rapid speciation and chromosomal evolution in mammals.” Proc Natl Acad Sci U S A 74(9): 3942-3946.

Böhne, A., F. Brunet, D. Galiana-Arnoux, C. Schultheis and J. N. Volff (2008). “Transposable elements as drivers of genomic and biological diversity in vertebrates.” Chromosome Res 16(1): 203-215.

Canapa, A., M. Barucca, M. A. Biscotti, M. Forconi and E. Olmo (2015). “Transposons, Genome Size, and Evolutionary Insights in Animals.” Cytogenet Genome Res 147(4): 217-239.

Castro-Insua, A., C. Gómez-Rodríguez, J. J. Wiens and A. Baselga (2018). “Climatic niche divergence drives patterns of diversification and richness among mammal families.” Sci Rep 8(1): 8781.

Cavalier-Smith, T. (1978). “Nuclear volume control by nucleoskeletal DNA, selection for cell volume and cell growth rate, and the solution of the DNA C-value paradox.” J Cell Sci 34: 247-278.

Chamberlain, S. A. and E. Szöcs (2013). “taxize: taxonomic search and retrieval in R.” F1000Res 2: 191.

Conger, A. D. and J. H. Clinton (1973). “Nuclear volumes, DNA contents, and radiosensitivity in whole-body-irradiated amphibians.” Radiat Res 54(1): 69-101.

Dores, R. M., C. Sollars, P. Danielson, J. Lee, J. Alrubaian and J. M. Joss (1999). “Cloning of a proopiomelanocortin cDNA from the pituitary of the Australian lungfish, Neoceratodus forsteri: analyzing trends in the organization of this prohormone precursor.” Gen Comp Endocrinol 116(3): 433-444.

Eastman, J. M. and A. Storfer (2011). “Correlations of life-history and distributional-range variation with salamander diversification rates: evidence for species selection.” Syst Biol 60(4): 503-518.

Einset, J. and A. R. Collins (2018). “Genome size and sensitivity to DNA damage by X-rays-plant comets tell the story.” Mutagenesis 33(1): 49-51.

Feder, J. L., R. Gejji, T. H. Powell and P. Nosil (2011). “Adaptive chromosomal divergence driven by mixed geographic mode of evolution.” Evolution 65(8): 2157-2170.

Feschotte, C. and E. J. Pritham (2007). “DNA transposons and the evolution of eukaryotic genomes.” Annu Rev Genet 41: 331-368.

Francis, D., M. S. Davies and P. W. Barlow (2008). “A strong nucleotypic effect on the cell cycle regardless of ploidy level.” Ann Bot 101(6): 747-757.

Gregory, T. (2004). “Macroevolution, Hierarchy Theory, and the C-value Enigma.” Paleobiology 30(2): 179-202.

Gregory, T. R. (2015). Animal Genome Size Database. http://www.genomesize.com.

Herrick, J. (2011). “Genetic variation and replication timing, or why is there late replicating DNA.” Evolution 65(11): 3031-3047.

Herrick, J. and A. Bensimon (2008). “Global regulation of genome duplication in eukaryotes: an overview from the epifluorescence microscope.” Chromosoma 117(3): 243-260.

Herrick, J. and B. Sclavi (2014). “A new look at genome size, evolutionary duration and genetic variation in salamanders.” Comptes Rendus Palevol 13(7): 611-615.

Hidalgo, O., J. Pellicer, M. Christenhusz, H. Schneider, A. R. Leitch and I. J. Leitch (2017). “Is There an Upper Limit to Genome Size?” Trends Plant Sci 22(7): 567-573.

Hooper, D. M. and T. D. Price (2015). “Rates of karyotypic evolution in Estrildid finches differ between island and continental clades.” Evolution 69(4): 890-903.

Kapusta, A., A. Suh and C. Feschotte (2017). “Dynamics of genome size evolution in birds and mammals.” Proc Natl Acad Sci U S A 114(8): E1460-E1469.

Karlin, A. A. and D. B. Means (1994). “Genetic variation in the aquatic salamander genus Amphiuma.” American Midland Naturalist 132 (1): 1-9

Kelly, L. J., S. Renny-Byfield, J. Pellicer, J. Macas, P. Novák, P. Neumann, M. A. Lysak, P. D. Day, M. Berger, M. F. Fay, R. A. Nichols, A. R. Leitch and I. J. Leitch (2015). “Analysis of the giant genomes of Fritillaria (Liliaceae) indicates that a lack of DNA removal characterizes extreme expansions in genome size.” New Phytol.

Knight, C. A., N. A. Molinari and D. A. Petrov (2005). “The large genome constraint hypothesis: evolution, ecology and phenotype.” Ann Bot 95(1): 177-190.

Kozak, K., D. Costantino, S. Lecaude, C. Sollars, P. Danielson and R. M. Dores (2005). “Analyzing the radiation of the melanocortins in amphibians: cloning of POMC cDNAs from the pituitary of the urodele amphibians, Amphiuma means and Necturus maculosus.” Peptides 26(10): 1920-1928.

Kozak, K. H. and J. J. Wiens (2010). “Accelerated rates of climatic-niche evolution underlie rapid species diversification.” Ecol Lett 13(11): 1378-1389.

Kozak, K. H. and J. J. Wiens (2016). “What explains patterns of species richness? The relative importance of climatic-niche evolution, morphological evolution, and ecological limits in salamanders.” Ecol Evol 6(16): 5940-5949.

Kraaijeveld, K. (2010). “Genome Size and Species Diversification.” Evolutionary Biology 37: 227-233.

Larson, A. (1981). “A reevaluation of the relationship between genome size and genetic variation.” The American Naturalist 118(1): 119-125.

Larson, A. (2011). “Developmental Correlates of Genome Size in Plethodontid Salamanders and Their Implications for Genome Evolution Author (s): Stanley K. Sessions and Allan Larson Reviewed work (s): Published by: Society for the Study of Evolution Stable URL: http://.” 41(6): 1239--1251.

Laurin, M., A. Canoville, M. Struble and V. Buffrénil (2015). “Early genome size increase in urodeles.” Comptes Rendus Palevol 15(1-2): 74-82.

Leaché, A. D., B. L. Banbury, C. W. Linkem and A. N. de Oca (2016). “Phylogenomics of a rapid radiation: is chromosomal evolution linked to increased diversification in north american spiny lizards (Genus Sceloporus)?” BMC Evol Biol 16: 63.

Lefébure, T., C. Morvan, F. Malard, C. François, L. Konecny-Dupré, L. Guéguen, M. Weiss-Gayet, A. Seguin-Orlando, L. Ermini, C. Sarkissian, N. P. Charrier, D. Eme, F. Mermillod-Blondin, L. Duret, C. Vieira, L. Orlando and C. J. Douady (2017). “Less effective selection leads to larger genomes.” Genome Res 27(6): 1016-1028.

Liedtke, H. C., D. J. Gower, M. Wilkinson and I. Gomez-Mestre (2018). “Macroevolutionary shift in the size of amphibian genomes and the role of life history and climate.” Nat Ecol Evol.

Lynch, M. and J. S. Conery (2003). “The origins of genome complexity.” Science 302(5649): 1401-1404.

Mank, J. E. and J. C. Avise (2006). “Cladogenetic correlates of genomic expansions in the recent evolution of actinopterygiian fishes.” Proc Biol Sci 273(1582): 33-38.

Marin, J. and S. Blair Hedges (2016). “Time best explains global variation in species richness of amphibians, birds and mammals” Journal of Biogeography 43: 1069-1079.

Marjanovic, D. and M. Laurin (2014). “An updated paleontological timetree of lissamphibians with comments on the anatomy of Jurassic crown-group salamanders (Urodela).” Historical Biology: An International Journal of Paleobiology 26(4): 535-550.

Martin, C. C. and R. Gordon (1995). “Differentiation trees, a junk DNA molecular clock, and the evolution of neoteny in salamanders.” J. Evol. Biol. 8(3): 339-354.

Martinez, P. A., U. P. Jacobina, R. V. Fernandes, C. Brito, C. Penone, T. F. Amado, C. R. Fonseca and C. J. Bidau (2017). “A comparative study on karyotypic diversification rate in mammals.” Heredity (Edinb) 118(4): 366-373.

Matsui, M., A. Tominaga, W. Z. Liu and T. Tanaka-Ueno (2008). “Reduced genetic variation in the Japanese giant salamander, Andrias japonicus (Amphibia: Caudata).” Mol Phylogenet Evol 49(1): 318-326.

McPeek, M. A. and J. M. Brown (2007). “Clade age and not diversification rate explains species richness among animal taxa.” Am Nat 169(4): E97-106.

Metcalfe, C. J. and D. Casane (2013). “Accommodating the load: The transposable element content of very large genomes.” Mob Genet Elements 3(2): e24775.

Mohlhenrich, E. R. and R. L. Mueller (2016). “Genetic drift and mutational hazard in the evolution of salamander genomic gigantism.” Evolution 70(12): 2865-2878.

Nevo, E. and A. Beiles (1991). “Genetic diversity and ecological heterogeneity in amphibian evolution.” Copeia 19(3): 565-592.

Oliver, M. J., D. Petrov, D. Ackerly, P. Falkowski and O. M. Schofield (2007). “The mode and tempo of genome size evolution in eukaryotes.” Genome Res 17(5): 594-601.

Olmo, E. (2006). “Genome size and evolutionary diversification in vertebrates.” Italian Journal of Zoology 73(2): 167 - 171.

Organ, C., M. Struble, A. Canoville, V. Bruffénil and M. Laurin (2016). “Macroevolution of genome size in sarcopterygians during the water–land transition.” Comptes Rendus Palevol 15(1-2): 65-73.

Organ, C. L., A. Canoville, R. R. Reisz and M. Laurin (2011). “Paleogenomic data suggest mammal-like genome size in the ancestral amniote and derived large genome size in amphibians.” J Evol Biol 24(2): 372-380.

Palazzo, A. F. and T. R. Gregory (2014). “The case for junk DNA.” PLoS Genet 10(5): e1004351.

Parker, E. and M. Kreitman (1982). “On the relationship between heterozygosity and DNA content.” The American Naturalist 119(5): 749-752

Pellicer, J., O. Hidalgo, S. Dodsworth and I. J. Leitch (2018). “Genome Size Diversity and Its Impact on the Evolution of Land Plants.” Genes (Basel) 9(2).

Pierce, A. and B. Mitton (1980). “The relationship between genome size and genetic variation.” The American Naturalist 116(6): 850-861.

Puttick, M. N., J. Clark and P. C. Donoghue (2015). “Size is not everything: rates of genome size evolution, not C-value, correlate with speciation in angiosperms.” Proc Biol Sci 282(1820): 20152289.

Pyron, R. A. and J. J. Wiens (2011). “A large-scale phylogeny of Amphibia including over 2800 species, and a revised classification of extant frogs, salamanders, and caecilians.” Mol Phylogenet Evol 61(2): 543-583.

Pyron, R. A. and J. J. Wiens (2013). “Large-scale phylogenetic analyses reveal the causes of high tropical amphibian diversity.” Proc Biol Sci 280(1770): 20131622.

Pyron, R. A. a. W. J. J. (2011). “A large-scale phylogeny of Amphibia including over 2800 species, and a revised classification of extant frogs, salamanders, and caecilians.” Molecular phylogenetics and evolution 61(2): 543--583.

Rabosky, D. L., G. J. Slater and M. E. Alfaro (2012). “Clade age and species richness are decoupled across the eukaryotic tree of life.” PLoS Biol 10(8): e1001381.

Schneider, H., H. Liu, J. Clark, O. Hidalgo, J. Pellicer, S. Zhang, L. J. Kelly, M. F. Fay and I. J. Leitch (2015). “Are the genomes of royal ferns really frozen in time? Evidence for coinciding genome stability and limited evolvability in the royal ferns.” New Phytol 207(1): 10-13.

Sessions, S. (2008). “Evolutionary cytogenetics in salamanders.” Chromosome Res. 16(1): 183-201.

Shaffer, B. H. and F. Breden (1989). “The relationship between allozyme variation and life history: non-transforming salamanders are less variable.” Copeia 4: 1016-1023.

Smith, E. M. a. G. T. R. (2009). “Patterns of genome size diversity in the ray-finned fishes.” Hydrobiologia 625(1): 1--25.

Smith, S. A. and B. C. O’Meara (2012). “treePL: divergence time estimation using penalized likelihood for large phylogenies.” Bioinformatics 28(20): 2689-2690.

Sparrow, A. H., C. H. Nauman, G. M. Donnelly, D. L. Willis and D. G. Baker (1970). “Radiosensitivities of selected amphibians in relation to their nuclear and chromosome volumes.” Radiat Res 42(2): 353-371.

Stadler, T., D. L. Rabosky, R. E. Ricklefs and F. Bokma (2014). “On age and species richness of higher taxa.” Am Nat 184(4): 447-455.

Sun, C., J. R. López Arriaza and R. L. Mueller (2012). “Slow DNA loss in the gigantic genomes of salamanders.” Genome Biol Evol 4(12): 1340-1348.

Van’t Hof, J. and A. H. Sparrow (1963). “A relationship between DNA content, nuclear volume, and minimum mitotic cycle time.” Proc Natl Acad Sci U S A 49: 897-902.

Vilenchik, M. M. and A. G. Knudson (2003). “Endogenous DNA double-strand breaks: production, fidelity of repair, and induction of cancer.” Proc Natl Acad Sci U S A 100(22): 12871-12876.

Vilenchik, M. M. and A. G. Knudson (2006). “Radiation dose-rate effects, endogenous DNA damage, and signaling resonance.” Proc Natl Acad Sci U S A 103(47): 17874-17879.

Vinogradov, A. E. (2003). “Selfish DNA is maladaptive: evidence from the plant Red List.” Trends Genet 19(11): 609-614.

Vinogradov, A. E. (2004). “Genome size and extinction risk in vertebrates.” Proc Biol Sci 271(1549): 1701-1705.

Wake, D. (2009). “What Salamanders have Taught Us about Evolution.” Ann. Rev. Ecol. Evol. Syst. 40: 333-352.

Wiens, J. J. (2017). “What explains patterns of biodiversity across the Tree of Life?: New research is revealing the causes of the dramatic variation in species numbers across branches of the Tree of Life.” Bioessays 39(3).

Wilson, A. C., G. L. Bush, S. M. Case and M. C. King (1975). “Social structuring of mammalian populations and rate of chromosomal evolution.” Proc Natl Acad Sci U S A 72(12): 5061-5065.

Wilson, A. C., V. M. Sarich and L. R. Maxson (1974). “The importance of gene rearrangement in evolution: evidence from studies on rates of chromosomal, protein, and anatomical evolution.” Proc Natl Acad Sci U S A 71(8): 3028-3030.

Zhang, H., S. Gao, M. J. Lercher, S. Hu and W. H. Chen (2012). “EvolView, an online tool for visualizing, annotating and managing phylogenetic trees.” Nucleic Acids Res 40(Web Server issue): W569-572.

